# SubcellulaRVis simplifies visualization of protein enrichment in subcellular compartments

**DOI:** 10.1101/2021.11.18.469118

**Authors:** Joanne Watson, Michael Smith, Chiara Francavilla, Jean-Marc Schwartz

## Abstract

High-throughput ‘omics methods result in lists of differentially regulated or expressed genes or proteins, whose function is generally studied through statistical methods such as enrichment analyses. One aspect of protein regulation is subcellular localization, which is crucial for their correct processing and function and can change in response to various cellular stimuli. Enrichment of proteins for subcellular compartments is often based on Gene Ontology Cellular Compartment annotations. Results of enrichment are typically visualized using bar-charts, however enrichment analyses can result in a long list of significant annotations which are highly specific, preventing researchers from gaining a broad understanding of the subcellular compartments their proteins of interest may be located in. Schematic visualization of known subcellular locations has become increasingly available for single proteins via the UniProt and COMPARTMENTS platforms. However, it is not currently available for a list of proteins (e.g. from the same experiment) or for visualizing the results of enrichment analyses.

To generate an easy-to-interpret visualization of protein subcellular localization after enrichment we developed the SubcellulaRVis web app, which visualizes the enrichment of subcellular locations of gene lists in an easy and impactful manner. SubcellulaRVis projects the results of enrichment analysis on a graphical representation of a eukaryotic cell. Implemented as a web app and an R package, this tool is user-friendly, provides exportable results in different formats, and can be used for gene lists derived from multiple organisms.

Here, we show the power of SubcellulaRVis to assign proteins to the correct subcellular compartment using gene list enriched in previously published spatial proteomics datasets. We envision SubcellulaRVis will be useful for cell biologists with limited bioinformatics expertise wanting to perform precise and quick enrichment analysis and immediate visualization of gene lists.

**Author Summary:** Cells contain intracellular compartments, such as membrane-bound organelles and the nucleus, and are surrounded by a plasma membrane. Proteins can be found in different cellular compartments; depending on the subcellular compartment they are localized to, they can be differentially regulated or perform location-specific functions. High-throughput ‘omics experiments result in a list of proteins or genes of interest; one way in which their functional role can be understood is through their subcellular localization, as deduced through statistical enrichment for Gene Ontology Cellular Component (GOCC) annotations or similar. We have designed a bioinformatic tool, named SubcellulaRVis, that compellingly visualizes the results of protein localization after enrichment and simplifies the results for a quick interpretation. We demonstrate that SubcellulaRVis precisely describes the subcellular localization of gene lists whose locations have been previously ascertained in publications. SubcellulaRVis can be accessed via the web or as a stand-alone app. SubcellulaRVis will be useful for experimental biologists with limited bioinformatics expertise analyzing data related to cellular signalling, protein (re)localization and regulation, and location-specific functional modules within the cell.

## Introduction

The localization of proteins within cells is critical for their processing, function, and regulation. Cellular homeostasis and response to environmental signals are also dependent on sequestering or dynamic re-localization of proteins in specific compartments [1]. Regardless of whether a protein is relocated in the cell or remains in a single compartment for the duration of its life, spatial regulation is critical for protein function. For example, upon translation, proteins bound for secretion will be trafficked through multiple compartments that compose the endomembrane system; contained within these compartments are “quality control” proteins that mediate correct folding or degradation for malformed proteins [2]. Cellular processes also occur in discrete, non-membrane bound locations, such as degradation within the proteasome, or transcriptional regulation in stress granules [3,4]. It is also increasingly appreciated that proteins which are dynamically recruited to different subcellular locations form functionally active modules that may be location specific, such as those that form and disperse at the lysosomal surface in response to growth factor or nutrient sensing [5].

Co-regulated proteins or genes are commonly identified using high-throughput ‘omics experiments, such as those based on transcriptomics, proteomics and phosphoproteomics. The regulated proteins or genes identified in these experiments are often assessed based on enrichment for annotations to particular biological characteristics or participation in biological pathways. Annotations based on subcellular localization are stored in the Gene Ontology aspect Cellular Component (GOCC) or the Jensen COMPARTMENTS database [6,7]. Enrichment for these annotations can provide initial indications of location-specific roles for a protein or gene list. Moreover, when analyzing spatially resolved data, annotations describing subcellular localization also provide an important data quality control. This is exemplified in the analysis of proteomics data that utilize proximity dependent biotinylation, in which bait proteins (known, characterized proteins tagged with a biotin ligase or peroxidase) covalently modify neighboring proteins (termed “prey”) through the addition of biotin. Isolating the biotinylated proteins generates a spatial interactome of the bait protein. Recently, BioID-based proximity-dependent biotinylation was used to generate cell-wide maps of proteins localized to particular organelles [8]. As demonstrated in this work, calculating enrichment for GOCC terms is a useful method for understanding whether the bait has been correctly tagged, in which case one would expect enrichment for the subcellular compartment the bait is commonly found.

Several web-based tools, such as EnrichR [9], can be used to perform enrichment analyses on protein or gene lists based on the GOCC or compartments databases. These analyses are accessible to biologists with minimal bioinformatics skills and are useful for understanding the subcellular compartments overrepresented in a list of proteins or genes of interest. However, a significant barrier in the interpretation of the results is that annotations are often highly specific, meaning it can be difficult to extract general trends from the potentially long list of returned annotations. For example, the enrichment for GOCC terms (calculated using EnrichR) on a list of GPI-anchored proteins (extracted from UniProt, Supplementary Table 1), results in a list of terms that are difficult to immediately generalize (Table 1). In contrast, certain cellular compartments are poorly characterized, for example the individual compartments of the endosomal system [10], and therefore analyses of proteins localized in such compartments may be noisy and incomplete.

**Table 1.**
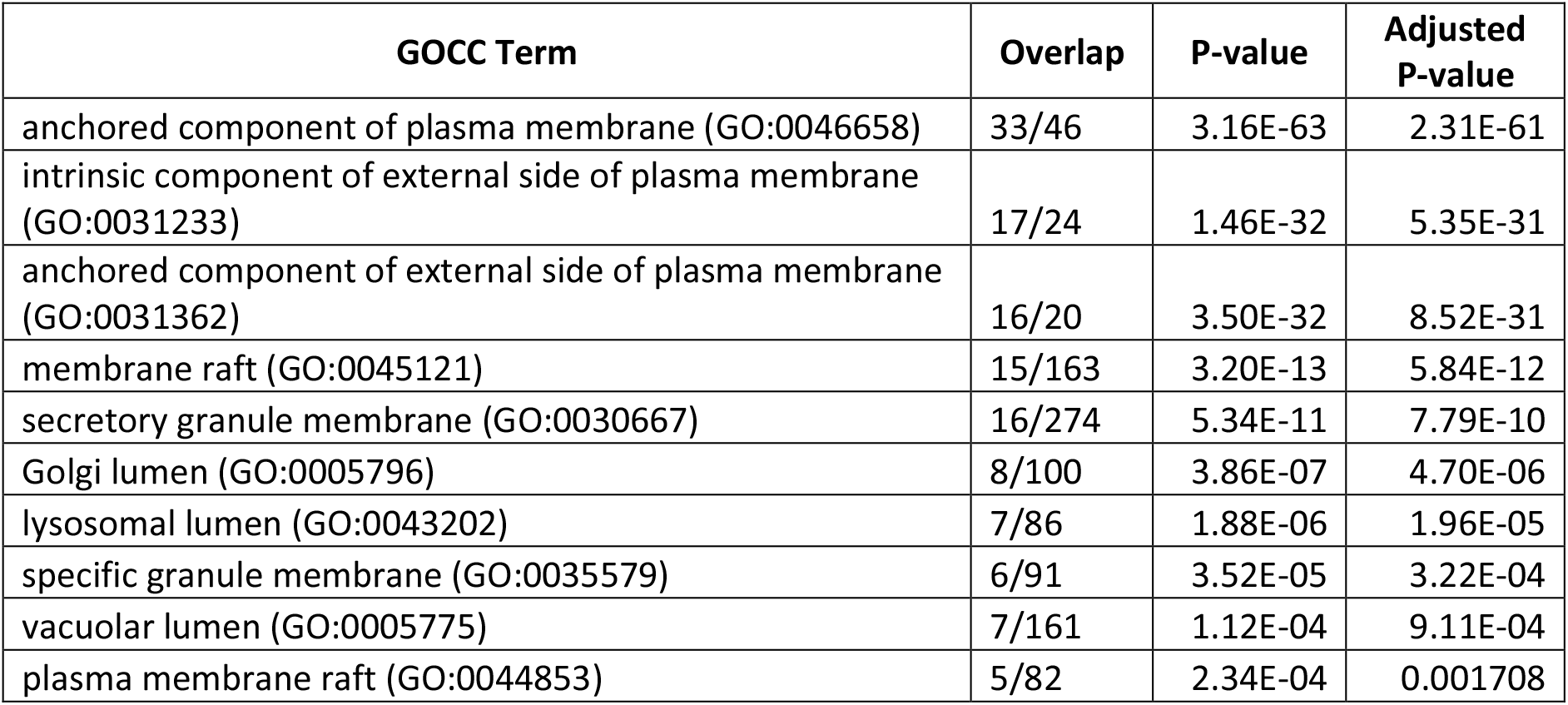
Results of GOCC enrichment from EnrichR on a list of 139 GPI-anchored proteins

We have implemented a Shiny web app, named SubcellulaRVis, that provides graphical visualization of subcellular compartment enrichment from a list of multiple genes or proteins. By performing the enrichment test using condensed annotations, so that there is a single term for each major intracellular compartment in the results, the analyses can be concisely visualized and interpreted. SubcellulaRVis represents an alternative to the visualization provided on the web interfaces of UniProt and COMPARTMENTS which only visualize the localization of a single protein [6,11]. A major novelty of our approach is the interactive visualization on a schematic of the eukaryotic cell of GOCC enrichment (Fig. 1), allowing for user-friendly interpretation, exploration, and presentation of the data. The visualization can be exported as a static image or as a table of the data behind the visualization. SubcellulaRVis provides a solution for non-bioinformaticians to investigate the subcellular localization of the proteins within a dataset of interest and standardizes the visualization of these results. The app is available at: http://phenome.manchester.ac.uk/subcellular/.

**Figure 1.**
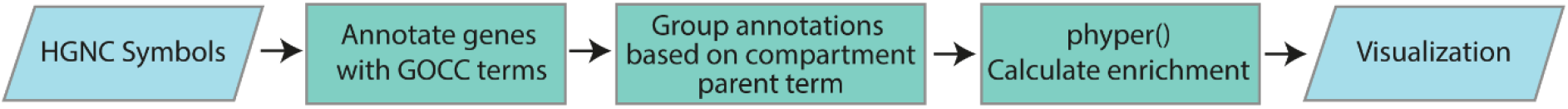
SubcellulaRVis workflow

## Design and Implementation

Our tool aims to simplify the interpretation of GOCC enrichment analyses through visualization of a standardized schematic of the cell. We selected subcellular compartments that are non-transient and common to eukaryotes. Though it is a non-exhaustive list, we anticipate it will meet the needs of the majority of cell biology researchers.

We first defined each cellular compartment visualized on the schematic (as shown in Figure 2) in respect to the GOCC terms that could describe it. We did this by utilizing the hierarchical structure of GO. The GO terms are hierarchically organized, with GO terms towards the “top” of the hierarchy being more general descriptors (e.g. GO:0005886, plasma membrane.) whilst terms towards the “bottom” of the hierarchy are more specific (e.g. GO:0005901, caveola) [12]. Each term in the GO hierarchy has parent terms (those that are higher in the hierarchy) and may also have child terms (those that are lower in the hierarchy). We identified the highest level GOCC term that would best describe each of the compartments visualized on the schematic (Table 1). Then, using the GO.db package in Bioconductor (version 3.13) [13], the child terms of the high-level GOCC terms were extracted and associated together. We call these grouped, compartment-specific terms the SubcellulaRVis compartment and use the high-level parent term to describe them, as shown in Table 1, hereafter. We extracted all the genes annotated to the terms in each SubcellulaRVis compartment using the species related AnnotationData libraries in Bioconductor (version 3.13), to create gene sets against which to calculate enrichment (Table 2). The annotation lists were generated for multiple species: *H. sapiens, M. musculus, D. melanogaster, S. cerevisiae, R. rattus,* and *X. laevis*.

**Figure 2.**
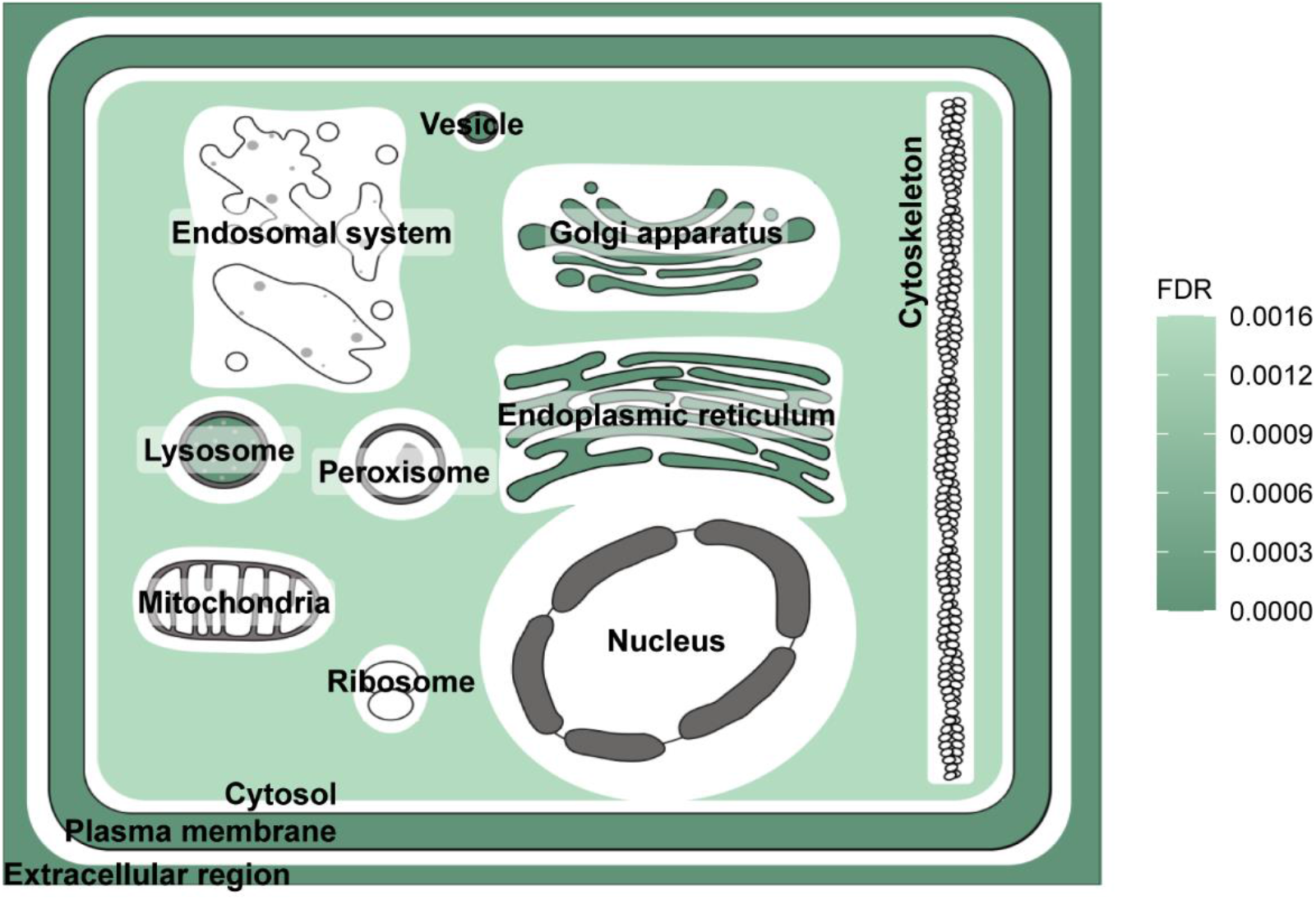
Visualization of enrichment results using SubcellulaRVis on a gene list of GPI anchored proteins (Supplementary Table 1) [11].

**Table 2.** Number of child terms for each general parent term selected.

| Compartment | GOID | Number of child terms | Number of genes annotated |  |  |  |  |  |
| --- | --- | --- | --- | --- | --- | --- | --- | --- |
|  |  |  | H. sapiens | M. musculus | D. melanogaster | S. cerevisiae | R. rattus | X. laevis |
| Cytoplasm | GO:0005737 | 1230 | 11847 | 11262 | 4700 | 4083 | 10436 | 4884 |
| Cytoskeleton | GO:0005856 | 273 | 2302 | 2213 | 523 | 255 | 2111 | 889 |
| Endoplasmic reticulum | GO:0005783 | 110 | 1961 | 1777 | 499 | 635 | 1700 | 685 |
| Endosome | GO:0005768 | 77 | 978 | 906 | 240 | 189 | 846 | 296 |
| Extracellular region | GO:0005576 | 94 | 4570 | 2697 | 1036 | 110 | 2072 | 1102 |
| Golgi apparatus | GO:0005794 | 47 | 1608 | 1468 | 379 | 285 | 1398 | 622 |
| Intracellular vesicle | GO:0097708 | 294 | 2405 | 1965 | 423 | 297 | 1848 | 555 |
| Lysosome | GO:0005764 | 38 | 697 | 472 | 106 | 6 | 456 | 145 |
| Mitochondrion | GO:0005739 | 96 | 1627 | 1852 | 863 | 1129 | 1570 | 650 |
| Nucleus | GO:0005634 | 492 | 8072 | 7127 | 2827 | 2302 | 6585 | 3673 |
| Peroxisome | GO:0005777 | 19 | 136 | 146 | 98 | 88 | 149 | 66 |
| Plasma membrane | GO:0005886 | 471 | 5726 | 5343 | 1383 | 507 | 5779 | 1878 |
| Ribosome | GO:0005840 | 20 | 243 | 231 | 285 | 267 | 295 | 239 |

Enrichment is calculated on the user supplied list of HUGO Gene Nomenclature Committee (HGNC) symbols (Figure 1). The genes are annotated to the GOCC terms based on the annotation lists generated for each species, and the annotations are then grouped based on the SubcellulaRVis compartment. A standard enrichment test is performed using the hypergeometric probability function, *phyper(),* in R, to calculate the enrichment for each SubcellulaRVis compartment in the user-supplied gene list, and corrected using the false discovery rate (FDR). Though it has limitations [14,15], the hypergeometric test was selected as values associated to the genes or proteins (e.g. expression or abundance values) do not need to be supplied in order to perform the test. To improve the accuracy of this calculation, the user can also input a background population of expressed genes or proteins in their sample, if known (as discussed in [16]). In the absence of the user-supplied background population list, the calculation is performed based on all the genes in the reference genomes (from the Bioconductor AnnotationData databases, as described above).

The SubcellulaRVis tool has been implemented as a Shiny web app (http://phenome.manchester.ac.uk/subcellular/) and a standalone app with R package (https://github.com/JoWatson2011/subcellularvis). The R package will be submitted to Bioconductor upon acceptance of the article. The visualization within the app has been implemented using ggplot2 and Plotly, allowing interactive exploration of the results when viewed on a web browser. All dependencies are described in the DESCRIPTION file of the R package (https://github.com/JoWatson2011/subcellularvis) and will be installed along with the package, as described in the README file.

## Results

### Features of SubcellulaRVis

The key feature of the SubcellulaRVis app is the visualization of GOCC enrichment results on a graphical representation of a eukaryotic cell (as seen in Figure 2), allowing for rapid and simple interpretation of the predominant localization of proteins of interest. By providing visualizing for a group of proteins, we provide an alternative to similar tools that visualize the localization of single proteins [6,11].

The app provides an interactive view of protein localization, and a static image of the plot can be exported; the characteristics of both the image and the plot (such as text size or color scale) can be chosen by the user. This is demonstrated by the test data, available in the app, which is a list of 139 proteins which are annotated as GPI-anchored in UniProt (Supplementary Table 1) [11]. Figure 2 visualizes the enrichment of these proteins for the membrane bound organelles of the secretory pathway, plasma membrane and for the extracellular space (Table 3). Cellular compartments in white are not enriched for the gene list.

**Table 3.** Enrichment results using SubcellulaRVis for gene list of 139 GPI-anchored proteins. Break in table indicates non-significant enrichment.

| Compartment | p-value | FDR | Number of genes in user list annotated to compartment |
| --- | --- | --- | --- |
| Plasma membrane | 4.70E-120 | 6.58E-119 | 137 |
| Extracellular region | 7.01E-83 | 9.11E-82 | 111 |
| Golgi apparatus | 1.15E-08 | 1.38E-07 | 21 |
| Intracellular vesicle | 1.34E-08 | 1.47E-07 | 26 |
| Endoplasmic reticulum | 3.61E-07 | 3.61E-06 | 21 |
| Lysosome | 4.84E-07 | 4.36E-06 | 12 |
| Vacuole | 1.97E-06 | 1.57E-05 | 12 |
| Cytoplasm | 0.000223 | 0.001559 | 57 |
| Endosome | 0.040296 | 0.241778 | 6 |
| Nucleus | 0.999908 | 1 | 10 |
| Cytoskeleton | 0.995838 | 1 | 1 |
| Mitochondrion | 0.995369 | 1 | 0 |
| Ribosome | 0.546016 | 1 | 0 |
| Peroxisome | 0.356874 | 1 | 0 |

Using the app or the associated R package, the user can view the results of the enrichment analysis graphically, as discussed above, or in tabular form. A standard enrichment including all the GOCC terms (rather than the summarized terms) can also be calculated and can be viewed in a separate tab on the app; also included is the SubcellulaRVis compartment each of these terms is assigned to, allowing the user full understanding of the way their data has been analyzed and visualized.

### Validation of GO Cellular Compartment Summarization

To assess the precision of the SubcellulaRVis compartment assignments, we validated the SubcellulaRVis tool using two previously published data sets describing subcellular localization of proteins. Both datasets utilized different spatial proteomics technologies in order to separate or label proteins found at different subcellular compartments.

The first dataset analyzed was from Go *et al.* [8] (from Supplementary table 8 of original publication). The authors used proximity dependent biotinylation to label proteins in the vicinity of different organellar markers which were tagged with the biotin ligase enzyme BioID. In total, 4424 proteins were identified and associated to different subcellular compartments. We calculated the enrichment using SubcellulaRVis of the proteins associated to the different organelles in the datasets, treating them as organelle-specific protein lists. The most significantly enriched SubcellulaRVis compartment for each of the twenty experimentally determined, organelle-associated protein lists were directly or closely matched (Table 4). For example, proteins experimentally associated to the cell junction, nucleolus and microtubules were enriched for the plasma membrane, nucleus, and cytoskeleton compartments, respectively by the SubcellulaRVis tool. The exception was the “Mitochondrial Membrane, Peroxisome”, as the proteins in this list were significantly enriched for the Cytoplasm. However, 21 / 172 proteins in this subset were annotated to both the mitochondria and the cytoplasm SubcellulaRVis compartments, and 6 / 172 were annotated to the peroxisome and the cytoplasm SubcellulaRVis compartments; none of the proteins in this subset were solely annotated to the mitochondria or the peroxisome, pointing to few location-specific markers for these compartments in the protein list. Moreover the ‘Mitochondrial Matrix’ and ‘Mitochondrial Inner Membrane/ Intermembrane Space’ showed significant enrichment for the mitochondria, indicating that our tool can associate known organelle markers to the correct compartment. This case study also provides a useful example of how analysis based on standardized, generic cellular compartments names, such as the SubcellulaRVis compartments, could be useful for comparison between different spatial studies.

**Table 4.** SubcellulaRVis enrichment results for each of the protein lists associated to cellular compartments by Go et al. [8]. The final column represents the overlap between the user-supplied gene list and genes annotated to the highest enriched SubcellulaRVis compartment.

| Experimentally determined Cellular Compartment | Highest enriched SubcellularVis compartment | FDR | No. User genes annotated to comp. / No. user genes |
| --- | --- | --- | --- |
| Cell junction | Plasma membrane | 1.88E-74 | 152/235 |
| Chromatin | Nucleus | 1.08E-216 | 352/383 |
| ER membrane | Endoplasmic reticulum | 7.11E-83 | 85/127 |
| Mitochondrial matrix | Mitochondrion | 8.78E-132 | 280/299 |
| Actin, cytosol | Cytoplasm | 6.19E-104 | 261/300 |
| ER lumen | Endoplasmic reticulum | 2.17E-127 | 122/165 |
| Endosome, lysosome | Endosome | 1.82E-97 | 88/173 |
| Nucleolus | Nucleus | 1.46E-85 | 146/164 |
| Nucleoplasm | Nucleus | 1.43E-131 | 222/247 |
| Nuclear body | Nucleus | 2.91E-179 | 278/295 |
| Plasma membrane | Plasma membrane | 1.04E-120 | 245/376 |
| Centrosome | Cytoskeleton | 7.48E-93 | 129/258 |
| Mitochondrial Membrane, Peroxisome | Cytoplasm | 3.78E-76 | 172/189 |
| Golgi | Golgi apparatus | 1.14E-100 | 96/146 |
| Nuclear outer membrane-ER membrane network | Endoplasmic reticulum | 5.58E-144 | 159/262 |
| Cytoplasmic RNP granule | Cytoplasm | 1.00E-43 | 119/143 |
| Microtubules | Cytoskeleton | 1.42E-86 | 87/115 |
| Mitochondrial inner membrane, Mitochondrial inter membrane space | Mitochondrion | 1.23E-134 | 106/122 |
| Misc. | Cytoplasm | 3.35E-86 | 234/279 |
| Early endosome, Recycling endosome | Intracellular vesicle | 1.03E-60 | 80/146 |

To assess the precision of the SubcellulaRVis tool for the analysis of a non-human data set we extracted the data from Nightingale et al. [17] (from Supplementary Table 3 of original publication) which used the hyperplexed Localisation of Organelle Proteins by Isotope Tagging (hyperLOPIT) method [18] to reconstruct spatial profiles of organelle proteomes in *Saccharomyces cerevisiae*. As with the previous dataset, SubcellulaRVis found the highest enrichment for the compartment most closely describing the experimentally associated cellular compartment, with the exception of the mitochondrion (Table 5). For example, the proteins lists associated to the endoplasmic reticulum and Golgi apparatus were associated with the respective SubcellulaRVis compartment. We concluded that SubcellulaRVis accurately replicates the protein localization determined in *S. cerevisiae*. Moreover, the SubcellulaRVis visualization (as demonstrated in Figure 1) would be an impactful way to display the enrichment results for each of the organelle-specific lists.

**Table 5.** SubcellulaRVis enrichment results for each of the protein lists associated to cellular compartments by Nightingale et al. [17]. The final column represents the overlap between the user-supplied gene list and genes annotated to the highest enriched SubcellulaRVis compartment.

| Experimentally determined Cellular Compartment | Highest enriched SubcellularVis compartment | FDR | No. User genes annotated to compartment / No. user genes |
| --- | --- | --- | --- |
| Cytosol | Cytoplasm | 9.47E-243 | 743/894 |
| ER | Endoplasmic reticulum | 4.55E-168 | 148/170 |
| Golgi | Golgi apparatus | 1.60E-111 | 80/109 |
| Mitochondrion | *No significant enrichment* | - | 12/186 |
| Nucleus | Nucleus | 1.55E-169 | 535/865 |
| Plasma membrane | Plasma membrane | 2.56E-118 | 160/401 |

## Availability and Future Directions

We have introduced an app named SubcellulaRVis, which provides easy visualization of the enriched subcellular locations from a user-supplied list of HGNC symbols. The visualization is more biologically insightful than a bar chart and will aid in efficient interpretation of the characteristics of gene lists and standardization of subcellular localization analyses. We anticipate our software will be particularly useful for experimental biologists with limited bioinformatics expertise analyzing data related to cellular signalling, protein (re)localization and regulation, and spatially resolved functional modules within the cell. For instance, our tool will provide impactful visualization of how protein localization varies upon two different extracellular perturbations, or for providing confirmation of correct tagging of proteins for spatial proteomics experiments.

SubcellulaRVis also provides an alternative view for the visualization of the endosomal system (Figure 3) which will be validated once appropriate data, such as spatial proteomics profiling of individual endosomal compartments (e.g. the early or recycling endosomes), becomes available. The endosomal system is notoriously hard to characterize [10], therefore our tool will be a valuable resource for distinguishing gene lists based on enrichment for different endosomal compartments. One potential use will be for researchers studying dynamic protein recycling to or from the plasma membrane.

**Figure 3.**
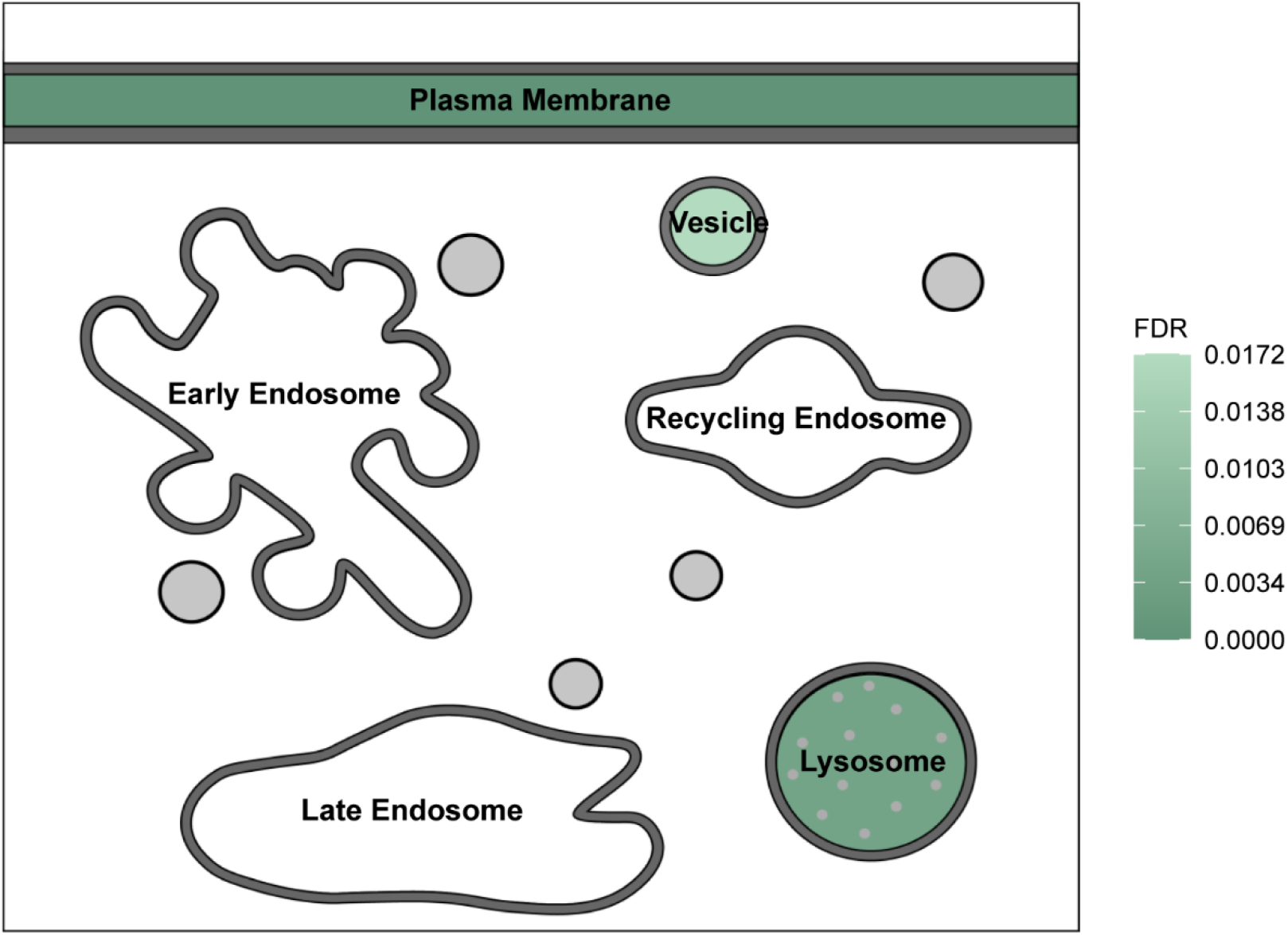
Visualization of enrichment results using SubcellulaRVis, using the trafficking-specific endosomal view, on a gene list of GPI anchored proteins (Supplementary Table 1 [11]).

The software can be installed at https://github.com/JoWatson2011/subcellularvis and the Shiny app is hosted at http://phenome.manchester.ac.uk/subcellular/. The R package will be submitted to Bioconductor upon acceptance of the article.

## Notes

http://phenome.manchester.ac.uk/subcellular/

### Competing Interest Statement

The authors have declared no competing interest.

